# A co-overexpression strategy allows effective protein-protein binding affinities to be assessed as a function of concentration within live cells

**DOI:** 10.64898/2026.07.20.739652

**Authors:** Samantha Stam

## Abstract

Weak, transient molecular interactions are ubiquitous amongst biological molecules. Regulatability by intracellular concentrations, localization, physical properties of the cytoplasm, and other factors is central to their importance. Here, we measure co-localization between fluorescently labeled proteins, Miro2 and β-Pix, to assess their effective binding affinity and its dependence on protein concentration. Co-localization between the two is quantifiable due to the mitochondrial localization of Miro2, which allows the otherwise cytosolic β-Pix to be drawn to mitochondria. Determining the fraction of overexpressed β-Pix co-localizing with mitochondria as a function of overexpressed Miro2 levels allows calculation of an effective dissociation constant from a mass-action binding model. Both the fit to the model and the effective dissociation constant for Miro2 and β-Pix binding depend on the concentration of β-Pix, which suggests that one or more model assumptions do not hold. Additional experiments reveal that competitive binding and/or β-Pix sequestration by Git1 are possible explanations for the β-Pix concentration dependence. Our modeling strategy is readily extendable to ectopic localization of proteins to mitochondria and other compartments for the assessment of co-localization with a binding partner. This reveals fundamental properties of protein-protein interactions and how they may be regulated by protein concentration and other intracellular conditions.

## INTRODUCTION

Weak interactions that occur between proteins and other biomolecules are a sufficiently central part of protein function and evolution that they have been designated as a fifth level of protein structure (*1*). The transient nature of these interactions is critical to maintaining dynamic, fluid structures such as lipid membranes with embedded proteins and liquid-liquid phase separated protein or nucleic acid droplets (*2, 3*). Reversibility of dynamic interactions allows a single protein to participate in multiple processes or to cooperate with other proteins to perform a function. These features may be favored by evolutionary fitness in higher organisms compared to strong interactions (*4*). However, higher affinity interactions also exist with dissociation constants in biomolecules often ranging from nanomolar to micromolar values (*5*). A robust way of measuring the effective affinity of binding inside of cells is needed to understand how and why these binding properties have evolved and how altering them influences cell physiology.

Existing techniques have shown that binding affinities of protein-protein interactions (PPIs) measured in cells may vary by orders of magnitude compared to their *in vitro* counterparts (*6*). The viscous and crowded physical environment of the cell has numerous effects on binding reaction equilibria and kinetics that may be dependent or independent of protein sequence (*7, 8*). Compartmentalization or confinement of proteins inside or outside of membrane-bound organelles, at contact sites between them, or inside of liquid-liquid phase separated droplets is another physical feature of the intracellular environment that can influence binding (*7*). Other factors that may alter binding affinities in cells compared to *in vitro* measurements include post translational modifications and competitive or cooperative binding from other proteins. Intracellular conditions may thus either produce different equilibrium dissociation constants than protein binding *in vitro* or may prevent measurement of a true dissociation constant (*5, 7, 8*). Despite the challenges for interpretation of PPI binding measurements in cells, detecting similar PPI dynamics between intracellular and *in vitro* conditions may be possible for some protein pairs (*9*). Thus, both equilibrium binding properties measured *in vitro* and intracellular conditions may influence the function and regulation of a PPI. Understanding the concentration dependence of a PPI in cells is critical to identifying any deviations from pairwise, equilibrium binding and how these deviations are regulated by intracellular conditions.

Previously established techniques for analyzing protein binding in cells are sufficiently quantitative to assess effective binding affinities but are limited to select proteins and concentrations. Forster resonance energy transfer (FRET) imposes limits on the protein concentration ranges due to background signal that depends on the protein stoichiometry (*10*) and must be corrected mathematically to detect PPIs (*11-13*). To additionally overcome this issue for the purpose of calculating dissociation constants, reference (*6*) varied the intracellular protein concentrations by altering cell volume via osmotic pressure. While this approach keeps the ratio of the protein concentrations within a binding pair the same, it simultaneously influences the concentrations of other intracellular factors and requires a model of how FRET varies with cell volume. In contrast, fluorescence correlation spectroscopy (FCS) does not have specific requirements for the ratio of proteins. However, FCS may only be suitable for select proteins and fluorescent tags and is susceptible to artifacts given its single-molecule sensitivity (*14, 15*). Thus, additional strategies are necessary for robust assessment of effective PPI strength at varying concentration and subcellular localizations that are relevant to conditions within cells.

Here, we present a method for using co-localization of two overexpressed fluorescently labeled proteins within a cell image to build a mass-action binding model at a range of protein concentrations. Similar to previous work (*16-21*), one protein is membrane-associated and draws the second, cytosolic protein to co-localize with the membrane. We advance these approaches to assess deviations from pairwise, mass-action binding. In our model, the fraction of the cytosolic protein that co-localizes with the membrane depends on the concentration of the first protein and an effective dissociation constant, the concept of which has been established in a previous FRET study (*22*). This analysis essentially reveals the dependence of a PPI on concentrations within individual cells as the concentrations are increased by overexpression. We use this model to infer that an association between the resident mitochondrial protein Miro2 (*23*) and the focal adhesion protein β-Pix (*24*) (Jodi Nunnari, personal communications) is limited by low endogenous concentrations natively present in the MDA-MB-231 cell line cytoplasm. The concentration limitation imposed by endogenous β-Pix deviates from the mass-action binding model. This suggests intriguing possibilities for regulation by effects including a competitive interaction or post-translational modifications (*25*) of β-Pix. Further examination reveals that competitive binding and/or sequestration by a second binding partner of β-Pix, Git1 (*26, 27*), could play roles in the β-Pix concentration dependence. Given its flexible application to a wide range of intracellular protein concentrations, our modeling approach is an important step toward understanding how PPIs depend on endogenous local or global concentrations and other effects present in the native intracellular environment.

## MATERIALS AND METHODS

### Plasmids and cell lines

The plasmids pGADT7-β-Pix isoform 4, pGBKT7-Miro2, pAC-BFP-Git1, pEGFP-C1, pAC-mCherry-C1, and kit components from the Matchmaker Gold yeast two-hybrid system (Takara Bio) were gifts from the Nunnari lab, UC-Davis. The MDA-MB-231 cell line was a gift from the Albeck lab at UC-Davis. As described below, cloning was conducted using these templates to make the pEGFP-β-Pix isoform 1 (referred to as EGFP-β-Pix in Results), pGADT7 or pEGFP-β-Pix mutants, pAC-mCherry-Miro2 (referred to as mCherry-Miro2), pAC-Git1 (referred to as unlabeled Git1), and pAC-SHD (referred to as unlabeled SHD) used for this study.

To make pGADT7-β-Pix isoform 1 from pGADT7-β-Pix isoform 4, the CH domain of β-Pix isoform 4 was removed via PCR opening of the plasmid. Prior to the PCR reaction, primers were phosphorylated with T4 Polynucleotide Kinase (T4 PNK, New England Biolabs (NEB)). An enzymatic digest with DpnI (NEB) was conducted to remove template plasmid from the PCR reaction, and re-ligation of the plasmid was done with Quick Ligase (NEB). Following these procedures, the c-terminus of β-Pix isoform 1 was ordered as a gBlock from Integrated DNA Technologies and swapped with that of isoform 4 by opening the vector with PCR and performing Gibson Assembly (Gibson Assembly Master Mix, NEB). Additional Gibson Assembly reactions were performed to add the β-Pix isoform 1 and Miro2 inserts from the yeast two-hybrid vectors pGADT7 and pGBKT7 to the mammalian cell imaging vectors pEGFP-C1 and pAC-mCherry-C1 respectively. After completing the pGADT7 and pEGFP vectors with β-Pix isoform 1 inserts, truncation mutants were made either by either the primer phosphorylation, PCR vector opening, DpnI digest, and re-ligation process described above or with Gibson Assembly. Similar procedures were used to make pAC-Git1 and pAC-SHD from the pAC-BFP-Git1 template. The EGFP-β-Pix mutant containing two point mutations used in Fig. 5 was made via two rounds of site-directed mutagenesis PCR performed with single primers and the enzymes T4 PNK, DpnI, and Taq Ligase (NEB). All PCR steps were done with Platinum SuperFi II polymerase (Invitrogen).

### Cell culture, transfection, and staining

MDA-MB-231 cells were cultured in DMEM media (Gibco) containing 10% fetal bovine serum (FBS, Gibco) and 100 U/mL penicillin-streptomycin (Gibco). The FBS was heat inactivated by incubation at 56° C for 30 minutes with occasional shaking. Cells were maintained in culture for less than 20 passages after thawing from frozen stocks.

Transfection of cells was conducted with Lipofectmine 2000 (Thermo Fisher) 24 to 36 hours before imaging. A total quantity of 1 µg of DNA using pcDNA3 as an empty vector in addition to the mCherry-Miro2, EGFP-β-Pix (wild type or mutants), and optional unlabeled Git1 or SHD plasmids. Transfections were done either for 4 hours in antibiotic-free minimum essential media (Opti-MEM, Gibco) or overnight in antibiotic-free full media. Immediately before imaging, cells were stained with 100 nM Mitotracker Deep Red FM (Invitrogen, catalog # M22426) in full media for 15 minutes. Destaining was conducted in full media for another 15 minutes before media replacement and imaging.

### Imaging and image analysis

Live-cell imaging was conducted at 60x magnification on a Zeiss LSM 980 equipped with an Airyscan 2 detector. For each cell analyzed, a z-series was completed for the entire cell volume with a spacing of 0.3 µm. Segmentation of mitochondria was done by manually determining a Mitotracker channel intensity threshold for each cell in ImageJ. Segmentation of the cells was done by manually outlining the EGFP-β-Pix channel excluding cell nuclei or vesicles in Matlab. Masks resulting from both segmentation procedures were used to select fluorescence intensities to sum or average over as described in Fig. 2A and the Results section.

## RESULTS

### Intracellular mass-action binding model suggests that an interaction between β-Pix and Miro2 is limited by endogenous Miro2 concentrations

We visualize an interaction between β-Pix and Miro2 previously validated by immunoprecipitation-mass spectrometry and yeast two-hybrid (Y2H) (Jodi Nunnari, personal communications) using images of MDA-MB-231 cells with overexpressed EGFP-β-Pix and mCherry-Miro2. Overexpression level is assessed via fluorescence intensity of either protein, and representative cells with varying overexpression are displayed in Fig. 1. The mCherry-Miro2 remains primarily localized to mitochondria at all overexpression levels as shown by the similarity between its images (middle column) and Mitotracker staining (last column), which is expected from previous studies (*23*). In contrast, EGFP-β-Pix localization, which is normally localized to focal adhesions or other membrane-associated puncta and the cytosol (*24, 28*) depends on overexpression level. When overexpression of EGFP-β-Pix, mCherry-Miro2, or both are sufficiently low, the EGFP-β-Pix in the first-column images remains mostly cytosolic with occasional intense localization near the cell periphery (Fig. 1 A-C, first column). At sufficiently high overexpression levels of both proteins, EGFP-β-Pix also localizes to mitochondria, which indicates that overexpressed β-Pix may be drawn to a non-native location by overexpressed Miro2 (Fig. 1D).

**Figure 1:**
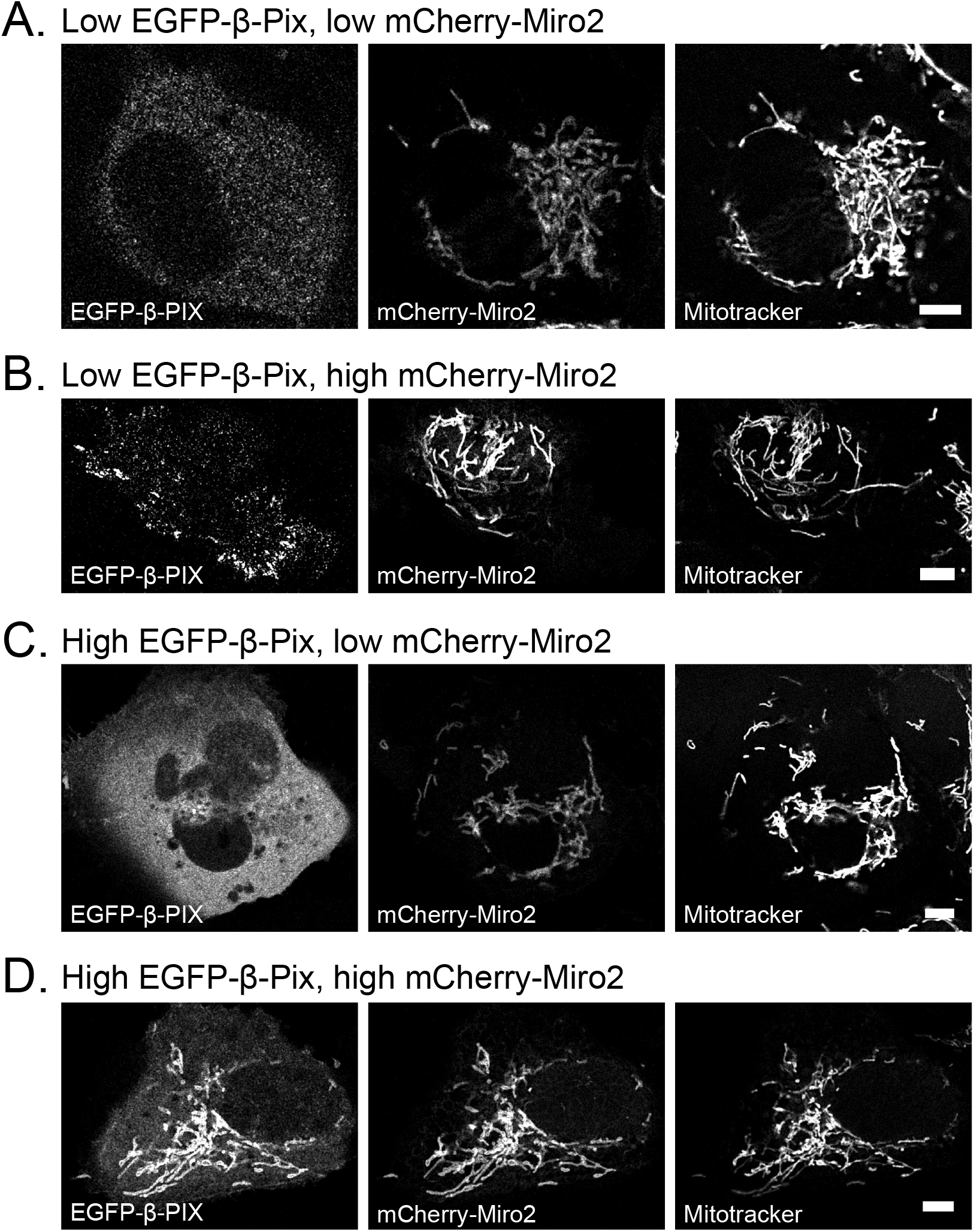
Overexpressed mCherry-Miro2 draws EGFP-β-Pix to mitochondria in a concentration-dependent manner. (A): MDA-MB-231 cell containing the lowest visible overexpression levels of EGFP-β-Pix (left panel) and mCherry-Miro2 (middle panel). Mitotracker staining (right panel) is used to visualize mitochondria. (B): Same as (A) but with higher mCherry-Miro2 overexpression. (C): Same as (A) but with higher EGFP-β-Pix overexpression. (D): Same as (A) but with higher overexpression of both proteins. Scale bars are 5 µm.

We examine the possibility that this effect could be used to understand the concentration dependence of the interaction. We rearrange the intensity data from each cell for fitting to a mass-action binding model:

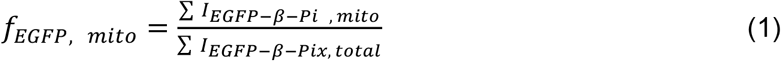

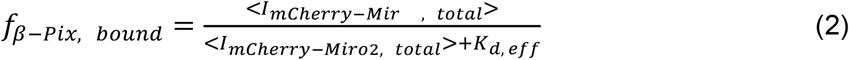

where *f*_EGFP, mito_ is the fraction of EGFP-β-Pix that overlaps with mitochondria determined experimentally, *f*_β-Pix bound_ is the fraction of intracellular β-Pix bound by Miro2, the *I* symbols represent fluorescence intensities of the species indicated in subscripts, and *K*_d,eff_, is an effective dissociation constant. These equations are used to examine the possibility that *f*_EGFP, mito_ calculated experimentally can serve as a proxy for *f*_β-Pix bound_. Notably, the measurement of a true dissociation constant using the form of equation (2) would assume that the majority of mCherry-Miro2 is unbound, the molecules are well-stirred and at thermodynamic equilibrium, and all molecules are equivalent, which are not necessarily true in our experiments (*5*).

To perform the calculation of Σ*I*_EGFP-β-Pix, total_ in equation (1), the EGFP-β-Pix channel is first used to segment entire cells in z-stack slices (Fig 2A, see Methods). Then, the intensity summation is conducted for all pixels within the segmented region. The same cell edge segmentation from the EGFP-β-Pix channel is used to obtain intracellular mCherry-Miro2 intensities for the calculation of <*I*_mCherry-Miro2, total_> in equation (2) (Fig. 2B). A third process in which the Mitotracker channel is used to segment mitochondria identifies the appropriate pixel intensities for Σ*I*_EGFP-β-Pix, mito_ in equation (1) (Fig. 2C). The resulting value of *f*_EGFP,mito_ is plotted against <*I*_mCherry-Miro2, total_> for each cell in Fig. 2D. A fit to equation (2) is performed by allowing *f*_EGFP,mito_ to serve as *f*_β-Pix,bound_ and using *K*_d,eff_ as the fitting parameter in Fig 2D. Consistent with mass-action binding, a rise followed by an apparent saturation of association between EGFP-β-Pix and mCherry-Miro2 at mitochondria with increasing mCherry-Miro2 is observed. Importantly, the interaction appears to be limited by endogenous Miro2 concentrations given the steeply rising slope of the curve near <*I*_mCherry-Miro2, total_> = 0.

**Figure 2:**
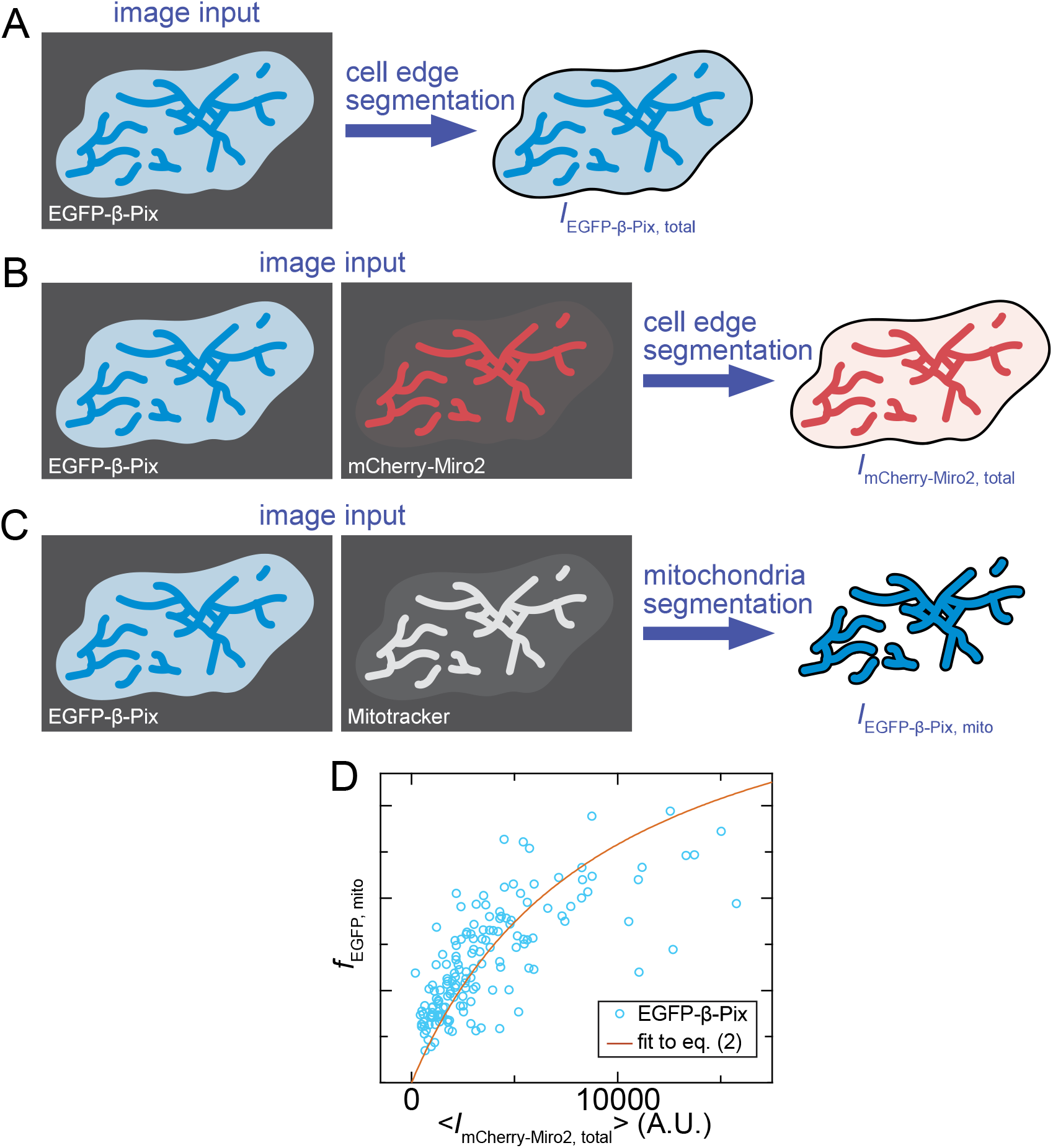
A mass-action binding model is fit to co-localization data of EGFP-β-Pix and mCherry-Miro2 at mitochondria. (A): Cell images in the EGFP-β-Pix channel (left) are used to segment the cell contents and determine the total EGFP-β-Pix intensity within a cell, *I*_EGFP-β-Pix, total_ (right). (B): The EGFP-β-Pix channel (left) is again used to segment cells and determine the portion of the mCherry-Miro2 channel (middle) to use for *I*_mCherry-Miro2, total_ (right). (C): The Mitotracker channel (middle) is used to segment mitochondria and select the portion of the EGFP-β-Pix channel (left) to use for *I*_EGFP-β-Pix, mito_ (right). (D): The resulting fraction of EGFP-β-Pix intensity co-localizing with segmented mitochondria, *f*_EGFP, mito_, against <*I*_mCherry-Miro2, total_> for individual cells. A fit to equation (2) (orange line) is performed by using *f*_EGFP, mito_ as *f*_β-Pix, bound_ and taking *K*_d, eff_ to be a fitting parameter.

### A background subtraction procedure reveals that the interaction is also limited by endogenous β-Pix levels

While the mass-action binding model in Fig. 2D resembles the trend of the data points, considerable deviations occur. Some of this variability may come from background overlap between mitochondria and EGFP-β-Pix that does not depend on the interaction. To estimate and correct for the background in each cell, we calculate the fraction of mitochondrial overlap of an EGFP-labeled β-Pix mutant, β-Pix_ΔGBD_, that lacks the minimal binding domain for the interaction (Git binding domain (GBD), Fig. S1). As expected for a construct that is mostly uniform in its distribution (Fig. 3A), this quantity does not vary with mCherry-Miro2 expression (Fig. S2A). Rather, *f*_EGFP_ for EGFP-β-Pix_ΔGBD_ rises linearly with the area fraction of Mitotracker in individual cells (*A*_mito_) with a slope of approximately one (Fig. 3B, open blue squares). Fitting a line to the data allows estimation of the background EGFP-β-Pix overlap with mitochondria as a function of *A*_mito_ (blue line, Fig. 3B).

**Figure 3:**
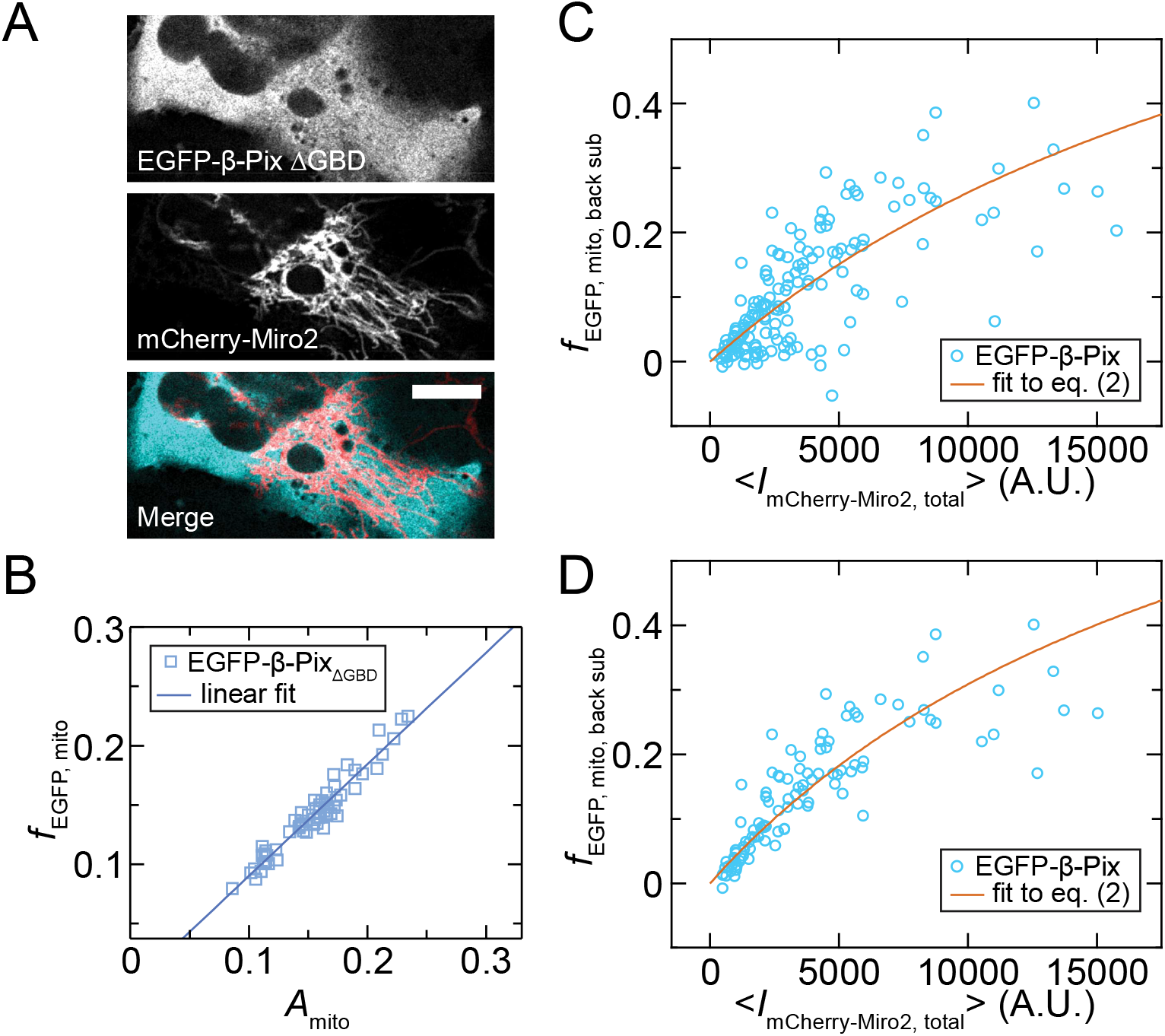
Background subtraction of EGFP-β-Pix overlapping with mitochondria indicates that the interaction is reduced at lower EGFP-β-Pix overexpression in a way that deviates from the mass-action binding model. (A): Images of cell overexpressing EGFP-β-Pix_ΔGBD_ (top) and mCherry-Miro2 (middle) at similarly high levels as in Fig. 1D. Merged image (bottom) shows mCherry-Miro2 in red and EGFP-β-Pix_ΔGBD_ in cyan. Scale bar is 10 µm. (B): Linear fit to *f*_EGFP, mito_ against total mitochondrial area, *A*_mito_, for the EGFP-β-Pix_ΔGBD_ mutant at all overexpression levels of it and mCherry-Miro2. (C): Background-subtracted fraction of wild-type EGFP-β-Pix intensity co-localizing with segmented mitochondria, *f*_EGFP, mito, back sub_ against <*I*_mCherry-Miro2, total_>. The fit line in (B) is used to calculate *b*_estimated_ in equation (3) for *f*_EGFP, mito, back sub_. The fit to equation (2) (orange line) is conducted using *f*_EGFP, mito, back sub_ as *f*_β-Pix, bound_. (D): Cells with <*I*_EGFP-β-Pix, total_> less than 1000 A.U. are removed before performing the same fitting procedure as in (C).

We further evaluate the appropriateness of the mutant for background subtraction by selecting cells overexpressing EGFP-β-Pix and mCherry-Miro2 at levels such that EGFP-β-Pix does not become concentrated on mitochondria. Similarity between the scaling of *f*_EGFP, mito_ with *A*_mito_ in these cells to those overexpressing EGFP-β-Pix_ΔGBD_ would indicate that the ΔGBD mutant is appropriate for background subtraction. Consistent with this, the fit line for *f*_EGFP_ versus *A*_mito_ in this subset of EGFP-β-Pix overexpressing cells is near that of the EGFP-β-Pix_ΔGBD_ overexpressing cells (Fig. S3).

Using this calculation, we replot the data from Fig. 2D with the estimated background, *b*_estimated_, subtracted from each data point:

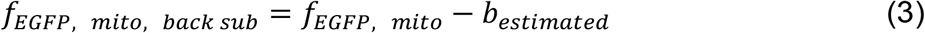

In Fig. 3C, the value of *b*_estimated_ for each cell is determined by inserting its mitochondrial area fraction into the EGFP-β-Pix_ΔGBD_ fit line from Fig. 3B. This procedure appears to reveal two populations of data points: one that outlines an apparent increasing trend with mCherry-Miro2 expression near or above the mass-action binding model fit line, and one that falls considerably below the fit line. These data further support the interaction being limited by intracellular Miro2 concentrations given that the data points at low overexpression of mCherry-Miro2 have a value of *f*_EGFP, mito, back sub_ near zero.

From the images of Fig. 1, we also expect low EGFP-β-Pix overexpression to limit the interaction. To reveal trends with changing EGFP-β-Pix concentration, we remove all data points from Fig. 3C with <*I*_EGFP-β-Pix, total_> less than 1000 A.U. and repeat the curve fit to equation (2) in Fig. 3D. This process causes a decrease of *K*_d,eff_ from 2.82 x 10^4^ to 2.24 x 10^4^ (Table 1). The model fit also improves from *r*^2^ = 0.56 to *r*^2^ = 0.71 (Table 1). These changes occur because the removed data points largely comprise the subset falling below the fit line in Fig. 2B. Together, these data reveal that endogenous concentrations of both Miro2 and β-Pix limit the binding interaction. For β-Pix, this occurs in a way that deviates from the mass-action binding model.

**Table 1:** Model fit information for different β-Pix overexpression conditions.

| Figure | Condition | $K_{d,eff}$ (A. U.) | $r^2$ |
| --- | --- | --- | --- |
| 2D | wild type (wt) β-Pix, without background subtraction | $9.37 \times 10^3$ | 0.40 |
| 3C | wt β-Pix, background subtraction | $2.82 \times 10^4$ | 0.56 |
| 3D | wt β-Pix, background subtraction, low β-Pix cells removed | $2.24 \times 10^4$ | 0.71 |
| 4C | β-Pix GBD, background subtraction | $1.24 \times 10^5$ | 0.59 |
| 4E | β-Pix GBD-CC, background subtraction | $3.47 \times 10^4$ | 0.60 |
| 5A | GBD-CC, background subtraction, low GBD-CC cells removed | $3.00 \times 10^4$ | 0.67 |
| 6B | β-Pix<sub>mono</sub>, background subtraction | $1.46 \times 10^5$ | 0.16 |
| S2B | β-Pix ΔGBD, background subtraction | $1.94 \times 10^6$ | 0.0215 |

### Multimerization-competent β-Pix construct containing the Git binding domain mimics binding of full-length β-Pix to Miro2

We examine the ability of our model to generate hypotheses explaining the observed concentration dependence of PPIs. To do this, we repeat our analysis on mutants of EGFP-β-Pix. Our choice of mutants is informed by yeast Y2H experiments indicating that the GBD (Fig. 4A, highlighted in dark green, (*27*)) is necessary and sufficient for the interaction between β-Pix and Miro2 (Fig. S1). Specifically, fragments of β-Pix containing various combinations of the SH3, DH, PH, and coiled-coil (CC) domains do not show activation in Y2H while the GBD alone activates (Fig. S1A). Consistent with this observation, the texture in the EGFP-GBD image of Fig. 4B appears to indicate weak co-localization with mCherry-Miro2 on mitochondria. Furthermore, the background-subtracted overlap with mitochondria of this construct shows an increase as a function of <*I*_mCherry-Miro2, total_> in Fig. 4C. The value of *K*_d,eff_ is raised from 2.82 x 10^4^ A.U. in Fig. 3C to 1.24 x 10^5^ A.U in Fig. 4C (Table 1). However, this value remains lower than that calculated using the same fitting procedure for the non-binding mutant, β-Pix ΔGBD, of 1.94 x 10^6^ A.U. (Fig. S2B, Table 1), which indicates that some co-localization of the β-Pix GBD is induced by mCherry-Miro2.

**Figure 4:**
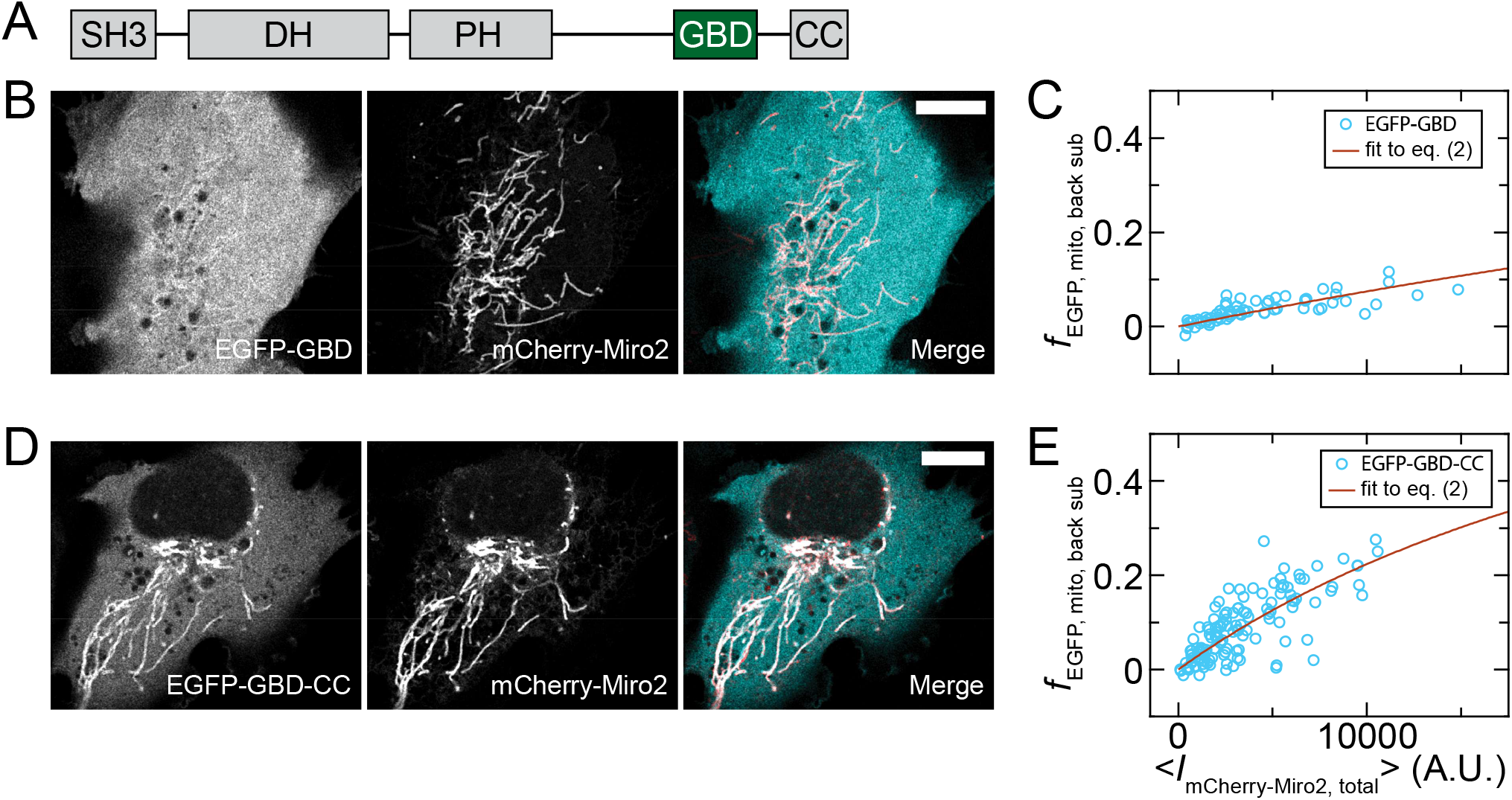
The CC multimerization domain enhances co-localization of the GBD with Miro2. (A): Schematic of the β-Pix structure including the minimal Miro2-binding domain, the GBD. (B): Images showing co-localization between overexpressed EGFP-GBD and mCherry-Miro2. Merged image shows mCherry-Miro2 in red and EGFP-GBD in cyan. (C): Values of *f*_EGFP, mito, back sub_ against <*I*_mCherry-Miro2, total_> for cells expressing EGFP-GBD. The fit to equation (2) (orange line) is conducted using *f*_EGFP, mito, back sub_ as *f*_β-Pix, bound_. (D) Images showing co-localization between overexpressed EGFP-GBD-CC and mCherry-Miro2. Merged image shows mCherry-Miro2 in red and EGFP-GBD-CC in cyan (E): Values of *f*_EGFP, mito, back sub_ against <*I*_mCherry-Miro2, total_> for cells expressing EGFP-GBD-CC. The fit to equation (2) (orange line) is conducted using *f*_EGFP, mito, back sub_ as *f*_β-Pix, bound_. Scale bars are 10 µm.

Reduced binding of the GBD alone could be influenced by the removal of the CC domain, which causes multimerization of β-Pix (*29, 30*) and influences association with its binding partner Git (*26, 30*). To examine the effect of this domain, we observe that restoring the CC appears to rescue co-localization of the EGFP-β-Pix fragment with mCherry-Miro2 at mitochondria (EGFP-GBD-CC, Fig. 4D). Quantifying this effect indicates that the construct increases localization to mitochondria relative to the GBD alone with a *K*_d,eff_ that remains only slightly higher than that of full-length EGFP-β-Pix (3.47 x 10^4^ A.U. compared to 2.82 x 10^4^, Fig. 4E, Table 1).

We use the GBD-CC truncation mutant to determine if these two domains are sufficient for the β-Pix concentration dependence observed in Fig. 3C-D. Like full-length β-Pix, omitting data points for which <*I*_EGFP-GBD-CC, total_> is less than 1000 A.U. results in removal of data points below the fit line (Fig. 5A compared to Fig. 4E). Repeating the curve fitting for a range of EGFP-β-Pix construct intensity cutoff values indicates that *K*_d,eff_ for both the full-length and GBD-CC constructs begins decreasing at values greater than 100 A.U. and then levels off for values greater than 500 A.U (Fig. 5B). The values of *K*_d,eff_ remain slightly higher for the EGFP-GBD-CC construct than for full-length EGFP-β-Pix at all cutoff intensities. In parallel, changes in the *r*^2^ trend with <*I*_EGFP-β-Pix, total_>_cutoff_ occur at similar values (Fig. 5C). These data suggest that the GBD-CC β-Pix truncation mutant is the minimal construct necessary to mimic binding of full-length β-Pix to Miro2.

**Figure 5:**
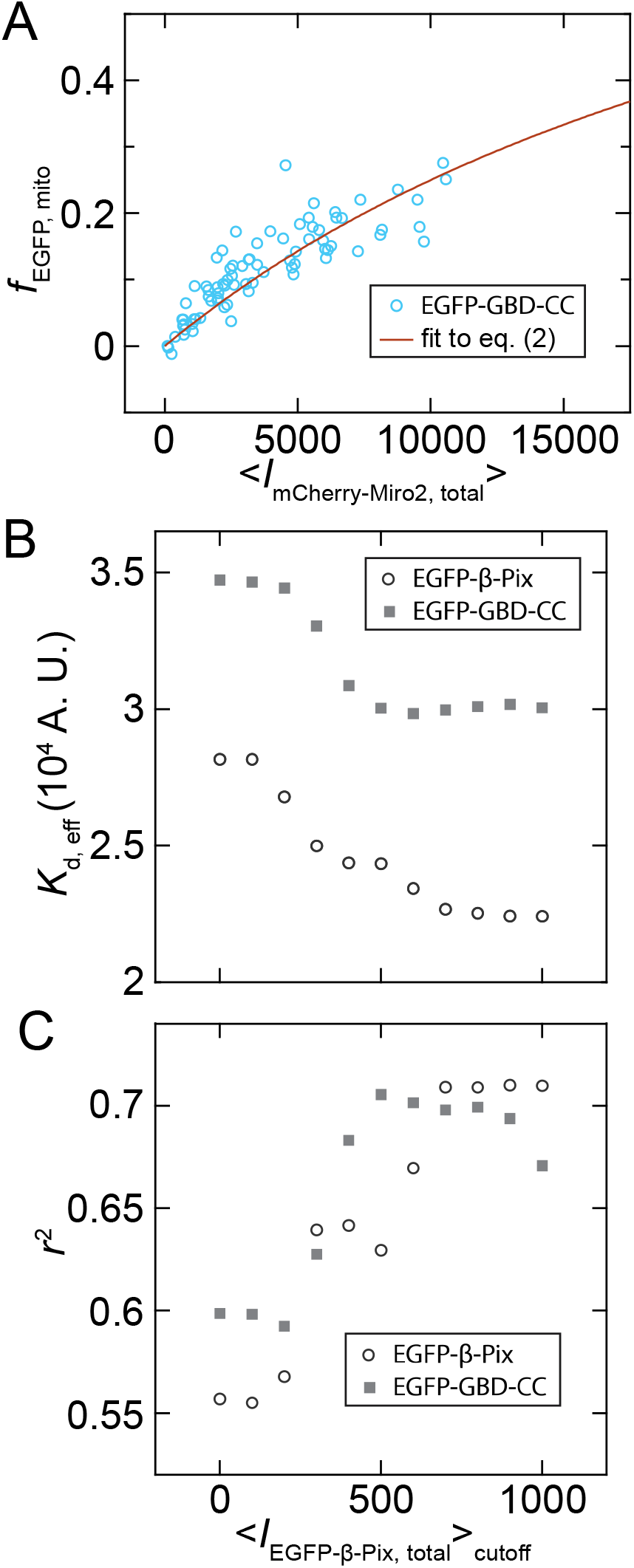
Both full-length EGFP-β-Pix and EGFP-GBD-CC show a sharp change in co-localization with mCherry-Miro2 at a similar overexpression level threshold. (A) : Cells with <*I*_EGFP-GBD-CC, total_> less than 1000 A.U. are removed before performing the same fitting procedure as in Fig. 4E. (B): Values of *K*_d, eff_ from fitting equation (2) to *f*_EGFP, mito, back sub_ versus <*I*_mCherry-Miro2, total_> for EGFP-β-Pix (open black circles) and EGFP-GBD-CC (filled gray squares). Cells with <*I*_EGFP-β-Pix, total_> less than <*I*_EGFP-β-Pix, total_>_cutoff_ are removed before fitting. (C): Values of *r*^2^ from the same curve fitting as in (B).

### Model indicates that a competitive interaction with Git1 and/or multimerization of β-Pix potentially influence the interaction between β-Pix and Miro2

The sufficiency of the GBD-CC construct for reduction in *f*_EGFP, mito, back sub_ values at low EGFP-β-Pix overexpression suggest potential explanations for the reduction. These include (1) competitive binding of the GBD by an endogenous factor or (2) sequestration of β-Pix into liquid-liquid phase separated domains containing Git (*26*), which might draw the β-Pix away from mitochondria. Either effect could potentially explain the relationship between *f*_EGFP, mito, back sub_ and EGFP-β-Pix overexpression by becoming saturated at high EGFP-β-Pix. We examine these possibilities with the model using co-overexpressed, unlabeled Git1 in addition to the EGFP-β-Pix and mCherry-Miro2. As would be expected for either a competitive interaction or competitive sequestration of EGFP-β-Pix by Git1, *f*_EGFP, mito, back sub_ is reduced at all values of <*I*_mCherry-Miro2, total_> (Fig. 6A, open green circles). This is consistent with images showing little co-localization of the EGFP-β-Pix and mCherry-Miro2 in Fig S4A. Co-overexpressing only the Spa2-homology domain (SHD) of Git1, which contains the binding site for β-Pix (*27*), is sufficient to cause similar reductions in *f*_EGFP, mito, back sub_ (Fig. 6A, open dark green squares) All data points in Fig 6A have an *I*_EGFP-β-Pix_ greater than 1000 A.U., so the reduction of binding at low EGFP-β-Pix is not expected to play a role.

**Figure 6:**
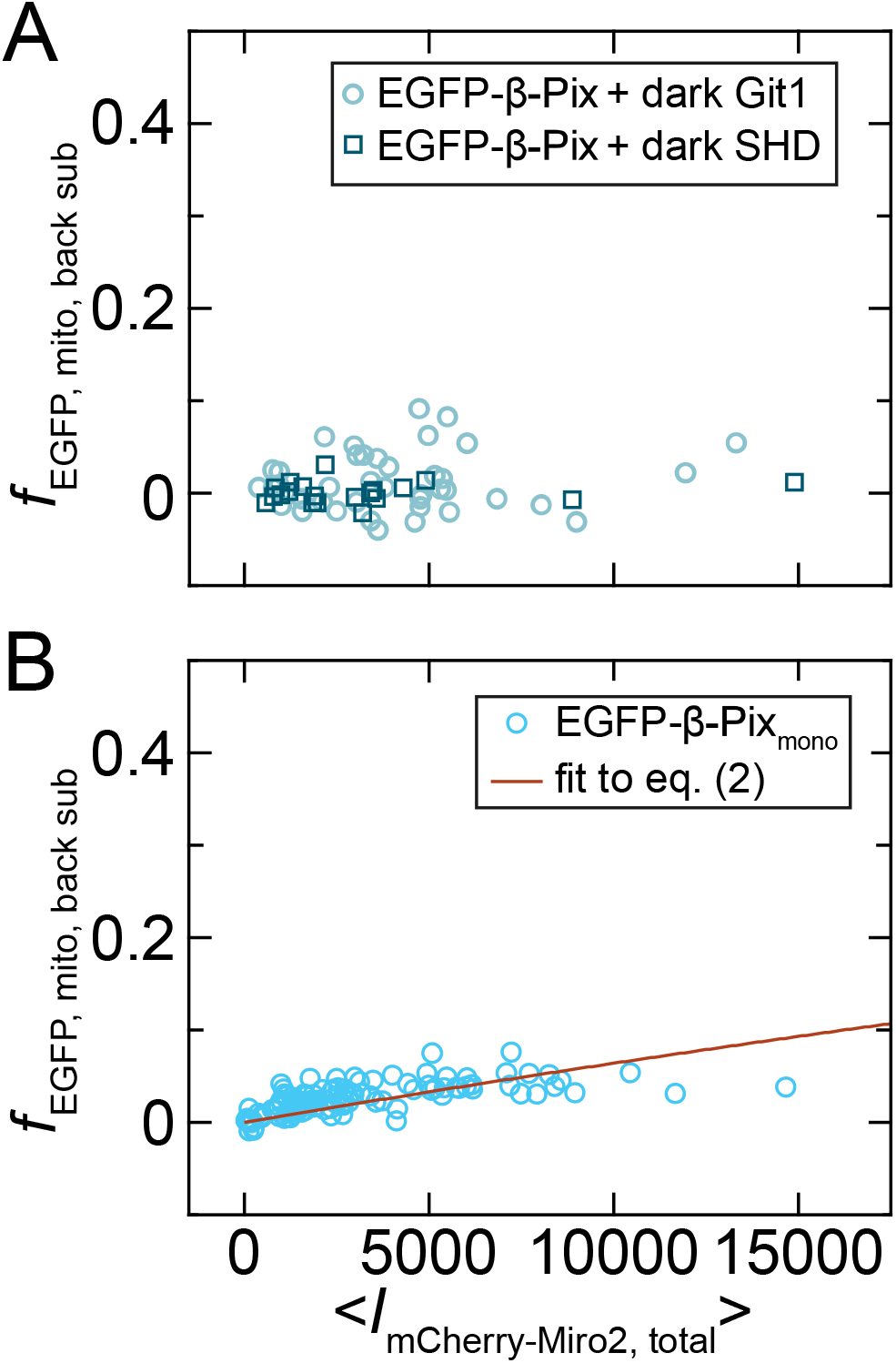
Model reveals possible regulatory mechanisms of the β-Pix-Miro2 interaction. (A): Background-subtracted *f*_EGFP, mito_ against <*I*_mCherry-Miro2, total_> for EGFP-β-Pix with co-overexpressed Git1 (open green circles) or the Git1 SHD (open dark green squares). (B): Background-subtracted *f*_EGFP, mito_ against <*I*_mCherry-Miro2, total_> for a monomeric β-Pix mutant, EGFP-β-Pix_mono_. The fit to equation (2) (orange line) is conducted using *f*_EGFP, mito, back sub_ as *f*_β-Pix, bound_.

Another observation from the mutants in Fig. 3 is that the GBD alone has reduced binding compared to either the full length β-Pix or GBD-CC constructs. To examine how multimerization via the CC domain influences effective binding affinity, we consider a mutant containing all domains of β-Pix and two point mutations in the CC that render it monomeric (β-Pix_mono_, (*26*)). These mutations also appear to reduce the co-localization with mCherry-Miro2 in images (Fig. S4B). Consistently, *K*_d,eff_ of the model fit is increased to 1.46 x 10^5^ A.U. (Fig. 6B, Table 1).

## DISCUSSION

Alterations in local or global protein concentration broadly underlie changes in cell function, morphology, and health. For example, low-affinity binding interactions may be enhanced by local compartmentalization and crowding (*7, 8*). Revealing these effects quantitatively will inform studies of membrane-bound or membraneless organelles, organelle-organelle contact sites, and protein aggregates that influence local protein concentrations. In addition to local concentration effects, global up or downregulation of protein concentrations across tissues may directly relate to the formation of bound complexes (*31*). Whether global or local protein concentration changes are at play, studying the role of concentration for PPIs requires assessment of effective binding affinity inside of the cells where the PPIs natively occur. Aside from the role of protein concentration effects in cell biology, this understanding will also be important for engineered systems that achieve local protein concentration increases inside of compartments or at interfaces with minimal components (*32-35*).

A specific example of potential regulation by local concentration revealed here is the effect of Git1 on β-Pix-Miro2 binding. While Git1 potentially competes with Miro2 for binding to β-Pix (Fig. 5) it may also locally concentrate β-Pix via liquid-liquid phase separation (*26*). An interesting possibility is that although Git1 inhibits the Miro2-β-Pix interaction when both proteins are at high concentration, Git1 could either enhance or spatially pattern the interaction at endogenous concentrations. Other questions of interest concern clarifying the effects of β-Pix multimerization via the CC domain. An intact multimerization domain both enhances phase separation of β-Pix into liquid droplets formed by Git1 (*26*) and enhances binding of β-Pix and Miro2 (Fig. 3C compared to Fig. 6B). Alternate isoforms of β-Pix that do not contain the CC domain are expressed in a tissue-specific manner (*29, 36*), which suggests another form of regulating the Miro2-β-Pix interaction via intracellular contents.

The modeling strategy we present (equation (1) and (2)) complements existing measurement techniques for binding affinities *in vitro* and *in vivo*. Intracellular environments may prevent determination of equilibrium dissociation constants by violating one or more of the following conditions: (1) the components are well-stirred, (2) all molecules of either interacting protein are equivalent, and (3) the concentration range is appropriate for the modeled relationship between the measured fraction of bound protein and the *K*_d_ to hold (*5*). Thus, purification and analysis of proteins *in vitro* will allow fuller interpretation of the binding curve obtained in cells. However, the deviations from equilibrium, mass-action binding evident in data from cells reveals intriguing possibilities. In comparison to the other *in vivo* methods for assessing binding affinities, FRET and FCS, this approach requires simple construct design and allows flexibility in the range of protein concentrations used. Additionally, given previous success in visualizing PPIs by ectopically localizing one of the interacting proteins to endosomal membranes (*19*), cytoskeletal filaments (*37*), or large artificial oligomers (*38, 39*), adapting our segmentation and fractional overlap technique to assess binding affinity with varying localization may be a future possibility. This approach would be difficult with FCS, which is nonideal for large objects of variable size (*14*). If one binding partner is ectopically located, competition with the endogenous protein in its native location will need to be considered during data interpretation. Appropriate background subtraction using a non-binding mutant or other cytosolic protein will also need to be customized for a given PPI. Unlike FRET and FCS, our technique is not amenable to two proteins that are both freely diffusing because one of the proteins must be anchored to a resolvable object.

A final application of our model is as a practical tool for determining cell lines that have stronger co-localization, and therefore more interacting molecules, at near-endogenous overexpression levels. This may be useful in selecting cell lines to use for downstream phenotypic analysis of a PPI even if the molecular mechanism for its relative enhancement in a cell line is unknown. Overall, our model will be a valuable tool in determining how PPI properties fit into the context of interaction networks in living cells.

## ACKNOWLEDGEMENTS

We thank Jodi Nunnari and members of the Nunnari lab, especially Yael Alon-Elbaz and Daniel Quinnell, for their helpful discussions and sharing of reagents. This work was supported by the National Institutes of Health NIGMS, project number 1R35GM141800-01.

