## Supplemental Methods and Figures for "A co-overexpression strategy allows effective protein-protein binding affinities to be assessed as a function of concentration within live cells"

Samantha Stam[1,2,*]

[1]Department of Molecular and Cellular Biology, University of California Davis, Davis, California and [2]James Franck Institute, University of Chicago, Chicago, Illinois.

### SUPPLEMENTARY MATERIALS AND METHODS

#### Yeast-two hybrid

The yeast strains Y2H Gold and Y187 from the Matchmaker Gold yeast two-hybrid kit (Takara Bio) were thawed from frozen stocks on yeast extract peptone dextrose (YPD, low ade) media plates. Following incubation for 2-3 days, overnight growth of single colonies was done in 5 mL liquid YPD cultures. Transformation-competent yeast cells for both strains were made by diluting these cultures in YPD and harvesting during logarithmic growth as determined by optical density (OD) measurements. The final transformation mixture consisted of cells pelleted from 5 mL of 0.6 OD culture, 33% polyethylene glycol, MW 3350 (Sigma), 0.1 M lithium acetate (Sigma), and 0.28 mg/mL salmon sperm DNA (Invitrogen). Vectors from the Matchmaker Gold kit, pGBKT7 and pGADT7, containing appropriate bait and prey genes were transformed and plated onto selective synthetic defined media (SD, SD -trp and SD -leu used for pGBKT7 and pGADT7 respectively) for 2-3 days. For additional selection of transformed cells, a second plating and incubation on selective media was performed. To obtain yeast containing both the bait and prey proteins, mating between singly transformed cells was conducted by mixing them on the surface of a YPD, low ade plate and incubating overnight. Two rounds of streaking on SD -leu -trp plates and incubation for 2-3 days were done to select for cells containing both plasmids. A final selection was conducted on SD -leu -trp -his plates to detect binding between the bait and prey. All yeast incubations were done at 30° C.

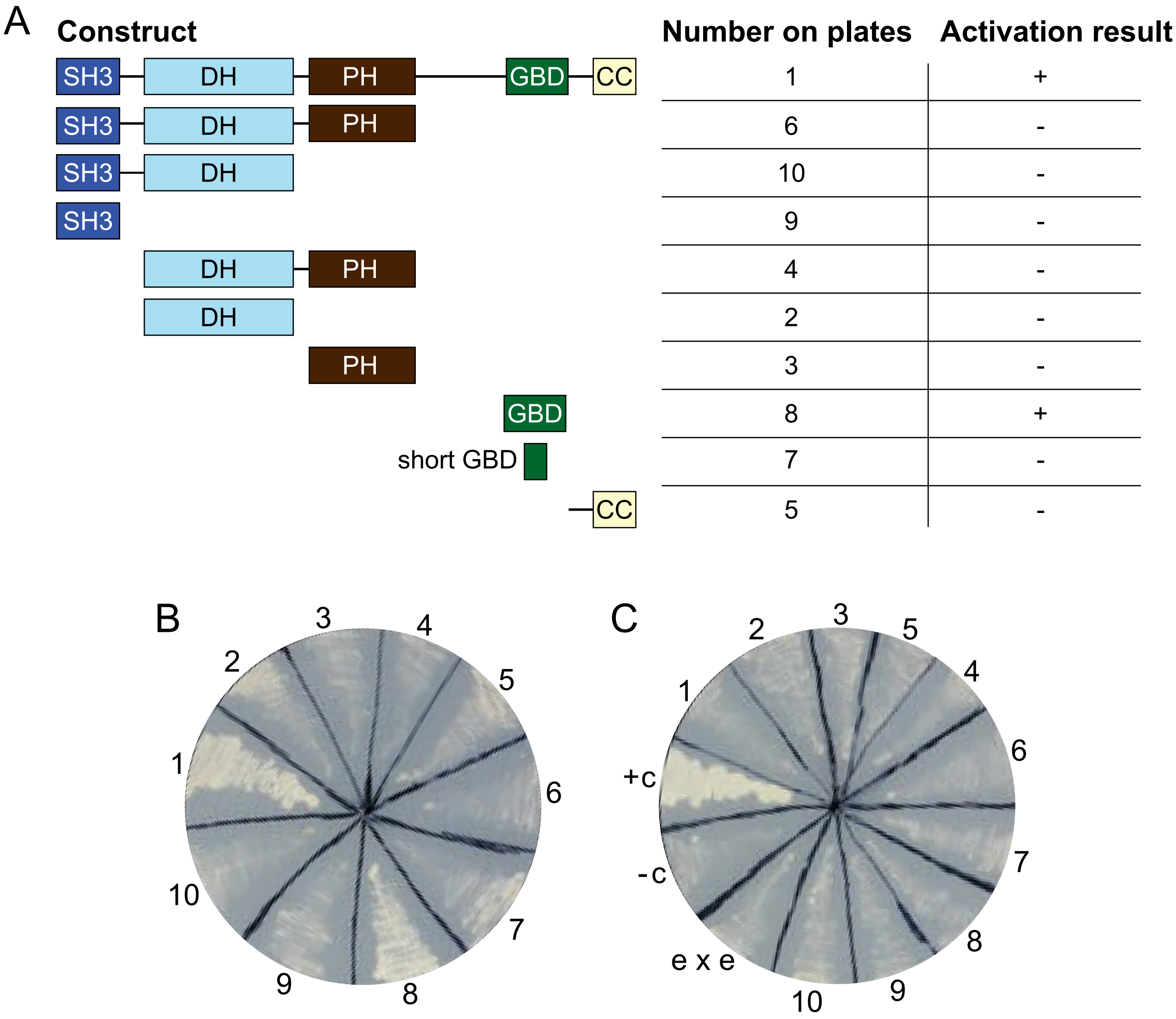


**Figure S1: Yeast two-hybrid (Y2H) indicates that the GBD of β-Pix is necessary and sufficient for binding to Miro2.** (A): Summary of activation results for yeast containing Miro2 and the indicated truncation of β-Pix. (B): Plate image of the results described in (A) with numbers indicating the β-Pix construct crossed with Miro2. (C): Control plate of yeast containing the numbered β-Pix construct and an empty vector. Additional controls include e x e, a cross containing both empty vectors, and +c or -c, the positive or negative controls from the Matchmaker Gold Y2H kit.

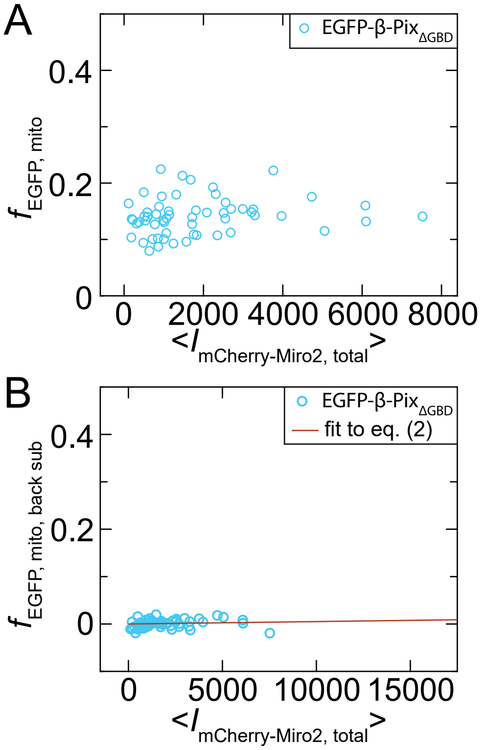


**Figure S2: Non-binding β-Pix mutant, β-Pix$_{\Delta GBD}$, does not show an increase in its mitochondrial overlap as Miro2 overexpression increases.** (A): Fraction of EGFP signal overlapping with mitochondria in cells overexpressing EGFP-β-Pix$_{\Delta GBD}$ as a function of average cellular mCherry-Miro2 intensity. (B): Same as (A) but with background subtraction using the curve in Fig. 3B and a fit to equation (2).

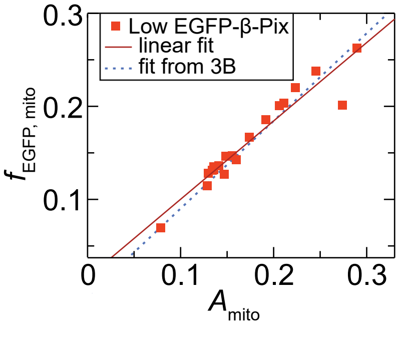


**Figure S3: At low overexpression levels, EGFP-β-Pix overlap with mitochondria shows similar scaling with mitochondrial area fraction as nonbinding mutant.** (A): Fraction of EGFP signal overlapping with mitochondria in cells overexpressing low levels of EGFP-β-Pix as a function of mitochondrial area fraction. The cells are also overexpressing mCherry-Miro2.

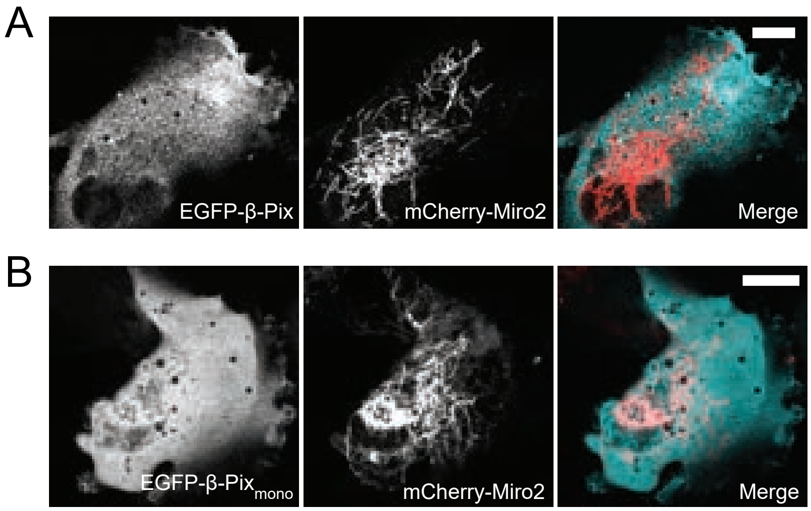


**Figure S4: Co-overexpression of Git1 and mutations of β-Pix rendering it monomeric both reduce binding to Miro2.** (A): EGFP-β-Pix, mCherry-Miro2, and merged image in a cell additionally co-overexpressing Git1. (B): EGFP-β-$Pix_{mono}$ mutant, mCherry-Miro2, and merged image.
